# Molecular Phenotypic Plasticity Informs Possible Adaptive Change of Triple-Negative Breast Cancer Cells *In Vivo*

**DOI:** 10.1101/2025.05.31.657204

**Authors:** Md. Iftehimul, Perpetua M. Muganda, Robert H. Newman, Checo J. Rorie, Scott H. Harrison, Muhammad T. Hossain, Misty D. Thomas, Joseph L. Graves, Dipongkor Saha

## Abstract

**Background and Objectives:** Cancer evolves via interconnected mechanisms, including changes in extrachromosomal DNA (ecDNA), genetic instability, and interactions with the tumor microenvironment (TME). These mechanisms allow for some clones to evolve metastatic traits, evade the immune system, and resist chemotherapy. However, how cancer cells evolve *in vivo* remains poorly understood. This study investigates the *in vivo* changes in gene expression of triple-negative breast cancer (TNBC) cells implanted in BALB/c mice.

**Methodology:** We analyzed RNA-seq data from 4T1 TNBC cells and tumors at different growth stages (1-, 3-, and 6-week) to identify differentially expressed genes, protein-protein interactions, and ecDNA alterations. We also assessed how ecDNA and genomic instability proteins interact with anti-TNBC drugs

**Results:** Our results reveal early transcriptional shifts within one week of tumor implantation, showing rapid acclimation. Changes in gene expression continued over time, with significant molecular reprogramming observed at six weeks under *in vivo* environmental pressures, including ecDNA alterations and immune evasion. The shift from the earlier generation (1 week) to the later generation (6 weeks) suggests cumulative alterations in key oncogenic pathways related to tumor progression. Additionally, we found that mutations in ecDNA and genomic instability proteins influence drug binding affinity, suggesting that adaptive changes may impact chemotherapy response.

**Conclusions and Implications:** This study provides new insights into how TNBC tumors may evolve over time and novel ecDNA-related mechanisms of possible tumor adaptation, highlighting potential biomarkers for tumor aggression and immune evasion, which could help develop more effective therapeutic strategies against TNBC.

## BACKGROUND and OBJECTIVES

Triple-negative breast cancer (TNBC) is an aggressive, metastatic, and heterogeneous subtype of breast cancer that lacks estrogen (ER), progesterone (PR), and human epidermal growth factor receptor 2 (HER2) receptors. The absence of these receptors makes it particularly challenging for TNBC to be treated with receptor-targeted therapies^[1]^. TNBC accounts for 10-15% of all breast cancer cases and is more prevalent in younger individuals, with variations in incidence across populations^[1]^. It has a higher recurrence rate, with nearly 34% of patients relapsing within five years, and a five-year survival rate for metastatic TNBC is as low as 12%^[2]^. Each year, around 200,000 new cases of TNBC are diagnosed worldwide, with approximately 40,000 deaths attributed to the disease^[3]^. By 2030, the global incidence of TNBC is projected to rise significantly due to increasing risk factors such as obesity and environmental influences^[4]^.

Cancer evolution is a dynamic process in which tumor cells first undergo several transcriptional modifications (acclimation) during proliferation and expansion, leading to intratumoral heterogeneity. This is followed by adaptation in which cells with advantageous mutations dominate, influencing tumor growth, metastasis, and therapy resistance^[5-8]^. Tumor evolution plays a crucial role in TNBC progression and poor prognosis^[9]^. TNBC progression involves multiple oncogenic pathways, including epithelial-to-mesenchymal transition (EMT), immune evasion, and alterations in DNA damage repair mechanisms^[10,11]^. A key concern in TNBC is its ability to evolve drug resistance. The process of drug resistance evolves first via acclimation that includes several mechanisms, including the upregulation of drug efflux pumps, alterations in apoptotic pathways, and adjustments to the tumor microenvironment (TME) that enable survival under therapeutic pressure^[12]^. This is followed by adaptation via genomic changes that encode the successful physiological changes, allowing tumor cells to proliferate in the host body^[13]^.

Chemotherapeutic agents like doxorubicin (DOX) and paclitaxel (PAC) are commonly used to treat TNBC^[14]^. DOX, an anthracycline, intercalates DNA, inhibits topoisomerase II, and alters the cellular redox environment via redox cycling in mitochondria^[15,16]^, while PAC, a taxane, stabilizes microtubules and disrupts mitotic progression^[17]^. However, TNBC frequently evolves resistance to these drugs through clonal selection and adaptive mutations. In tumors, selective pressures imposed by chemotherapy and the immune system drive the evolution of resistant clones^[18]^. This ongoing battle between therapy and tumor adaption necessitates the development of novel therapeutic strategies that can preemptively or concurrently target resistance mechanisms. Among the factors influencing tumor acclimation and subsequent adaptation, extrachromosomal DNA (ecDNA) plays a crucial role in tumor evolution^[19]^ by facilitating rapid genomic modifications, promoting tumor cell proliferation, and enhancing treatment resistance^[20,21]^. Understanding how ecDNA contributes to TNBC evolution and drug resistance may provide valuable insights into overcoming therapeutic challenges.

The 4T1 cell line, a highly metastatic murine TNBC model, is widely used to study TNBC progression and resistance^[22]^. This study investigates how 4T1 cells undergo early acclimation and subsequent changes in an *in vivo* environment and how these could potentially contribute to drug resistance. Using the RNA-seq datasets from NCBI, we tracked the transcriptional trajectory of 4T1 cells across three stages of *in vivo* tumor growth: early-stage (1-week growth), intermediate-stage (3-week growth), and late-stage (6-week growth). We examined ecDNA alterations across different stages of tumor growth and their impact on drug-binding affinity. As the tumor cells grow under environmental pressure, mutator alleles may appear, driving increased mutation rates and drug resistance. Thus, we also assessed the interaction of anti-TNBC drugs with potential mutator alleles to understand their role in therapy resistance.

## METHODOLOGY

### Datasets acquisition and statistical analysis of differentially expressed genes (DEGs)

To determine acclimatory changes of 4T1 tumors over time *in vivo*, we extracted RNA-seq datasets (originating from 4T1 cells and 1-week-, 3-week-, and 6-week-old 4T1 tumors) from the NCBI database (*Table S1*) and aligned the reads to the GRCm39 genome using HISAT2^[23]^. We quantified gene-level read counts using FeatureCounts^[24]^, normalized them across samples using DESeq^[25]^, and analyzed DEGs. In this study, genes with absolute p-value <0.05 and absolute Log2FC >1 were considered to be potential DEGs. Additionally, we used Log2FC ≥1 and Log2FC ≤ −1 criteria to explore up-and down-regulated genes, respectively, and further calculated for adjusted p-values. DEGs were then imported into R Studio to generate the volcano plots, principal component analysis (PCA), and heatmaps^[26]^. Additionally, Jvenn was utilized to identify DEGs shared across all groups^[27]^.

### Enrichment analysis of shared DEGs

To further explore the phenotypic plasticity of signaling pathways during carcinogenesis, we used shared DEGs to construct a protein-protein interaction (PPI) network and KEGG pathways. The PPI network was constructed using the STRING database^[28]^ with the following criteria: the selected species was *Homo sapiens*, and a reliability threshold of a medium confidence score of 0.4 was chosen to balance coverage and accuracy, ensuring a well-connected network with reliable interactions while minimizing false-positive interactions. Furthermore, PPI information was analyzed to retrieve the topological properties of nodes in the core network, such as betweenness centrality (BC) and closeness centrality (CC) values, using Cytoscape v3.7.2^[29]^. The cytoHubba plugin was used to determine the top connective nodes, while module clustering within the PPI network was performed using the MCODE plugin. To identify relevant pathways, we conducted a KEGG pathway enrichment analysis on all possible common genes using the DAVID database^[30]^, retaining only those pathways with a significance threshold of p < 0.05. Similarly, ShinyGo 0.80 was applied to illustrate the enrichment analysis of KEGG results in a dot plot^[31]^.

### Interaction of ecDNA-related oncoproteins and anti-TNBC drugs

The three-dimensional structures of wild-type and mutated ecDNA-related oncoproteins were obtained from the Protein Data Bank. The protein structures were selected based on their crystallographic data, with metals, co-factors, waters, and side chains excluded. The active residues of these proteins were identified using the COACH-D algorithm. To enhance the specificity of the screening, chain A was selected and prepared by removing unwanted ions, heteroatoms, ligands, and water molecules. Additionally, hydrogen atoms were added to the proteins to ensure proper functioning during the screening process. All protein preparation steps were performed using the UCSF Chimera 1.17.3 platform^[32]^.

The molecular screening study was conducted using the PyRx 0.8 package^[33]^, a computational molecular docking toolkit. A grid box was created within the active site of the core protein, with dimensions x = 28.33, y = 35.10, and z = 39.48. The grid center was positioned at coordinates x = 26.72, y= −10.14, and z = 1.45^[34]^. The grid box position was kept constant for each selected core protein to maintain a consistent screening environment. AutoDock Vina was run via a batch script, which sequentially executed commands to operate the software^[35]^. Following the docking process, multiple models (usually between 1 and 9) were generated for each receptor-ligand interaction. The model with the highest binding free energy and the lowest Root Mean Square Deviation (RMSD) score was identified as the optimal one^[36]^. Hydrophobic and hydrogen bond interactions were further highlighted in the molecular interaction analysis using BIOVIA Discovery Studio^[37]^.

## RESULTS

### Acclimatory transcriptional shift in 4T1 cells: Insights from DEG analysis

Cancer cells first undergo rapid acclimation followed by evolutionary changes, driven by mutation and selection due to internal and external pressures, which results in adaptation displayed at both molecular and cellular levels^[38,39]^. DEG analysis provides a crucial tool to uncover evolutionary transcriptional changes across generations of tumor cell growth, shedding light on key drivers of drug resistance^[40]^. By analyzing gene expression patterns at different time points—1-week (1w), 3-week (3w), and 6-week (6w)—we can trace the phenotypic trajectory of tumor cells and identify genes that are significantly upregulated or downregulated, providing a vision of possible future adaptation^[13,41]^.

As indicated in *Fig. 1A-C*, during the initial acclimation phase (1w) in an *in vivo* model system, 4T1 tumor cells undergo rapid transcriptional changes, as 14,571 genes are differentially expressed, with 6,507 genes upregulated. As the 4T1 cells progress to the intermediate stage (3w), molecular acclimations persist, albeit at a slower rate, with 6,799 genes upregulated. By the late stage (6w), 4T1 cells show an even more transcriptionally changed phenotype that may contribute to enhanced survival mechanisms, increased drug resistance, and extensive genetic and epigenetic alterations^[42]^. At this point, only 8,925 genes remain differentially expressed, with 3,973 genes upregulated, indicating that roughly 60% of the previously acclimatized genes have stabilized under *in vivo* environmental pressures, possibly due to selection pressure from the host immune responses^[43,44]^. Similarly, as shown in the heatmap analysis, we observed transcriptional shifts at different stages of 4T1 tumor growth (from week 1 to week 6) compared to the baseline 4T1 cells (*Fig. 1D*).

**Figure 1:**
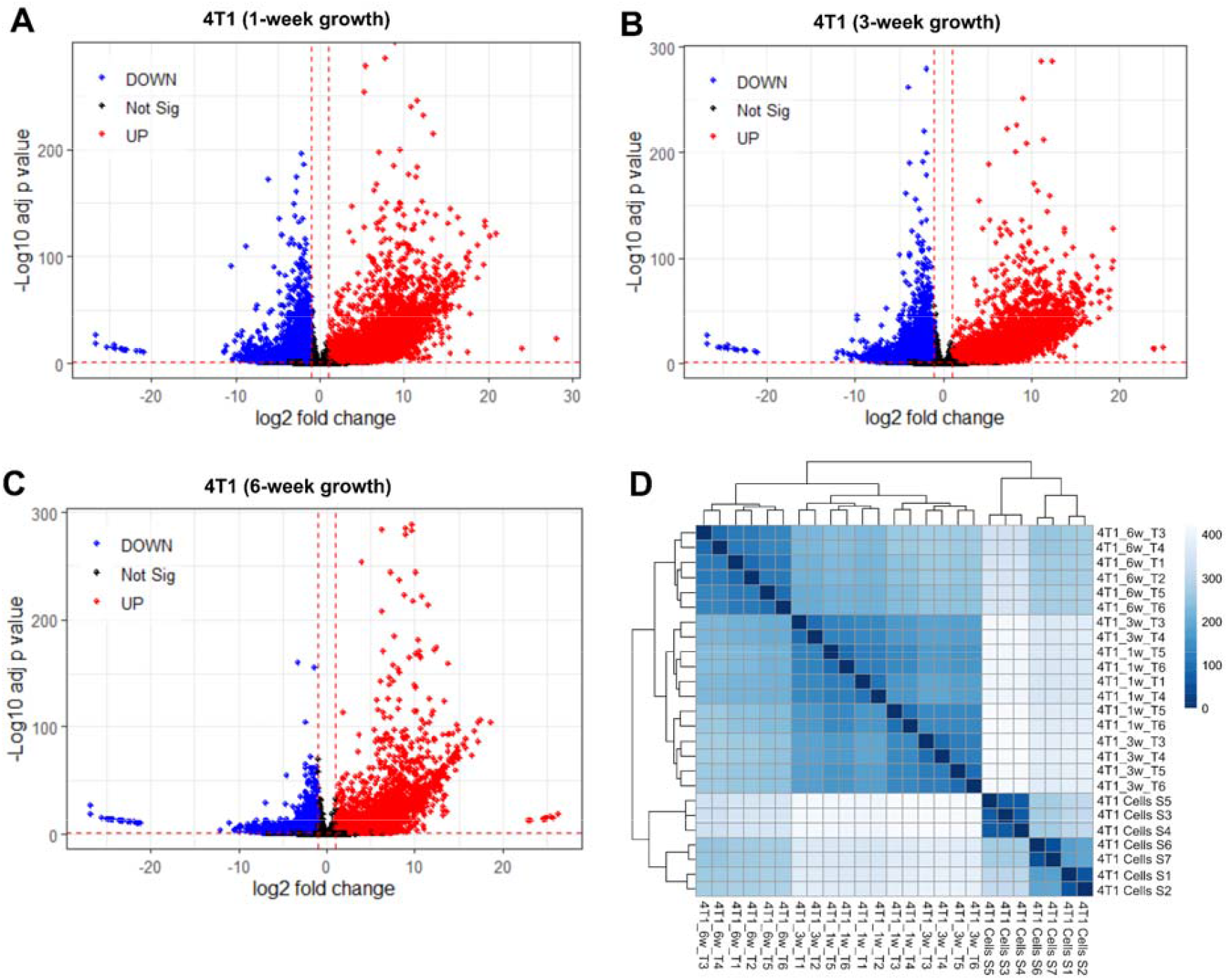
**(A-C)** Volcano plot at different stages of 4T1 tumor cell growth. Using 4T1 cell RNA-seq as a baseline dataset, these plots show gene expression changes at (A) 1-week (14,571 DEGs), (B) 3-week (15,036 DEGs), and (C) 6-week (8,925 DEGs) post-tumor implantation. The x-axis (log2 fold change) indicates the magnitude and direction of expression changes, with positive values denoting higher expression and negative values denoting lower expression. The y-axis (labeled as “Log10 adj. p-value”) reflects statistical significance, where higher values (e.g., 0) correspond to lowest adjusted p-values (e.g., 1e-200), highlighting genes with highly confident differential expression. The plot reveals numerous genes with large fold changes (far left/right on the x-axis) and extreme significance (top of the y-axis), suggesting substantial molecular shifts during tumor progression. DOWN – downregulation; Not Sig – not significant; UP – upregulation. **(D)** Sample-to-sample distance heatmap to understand the gene expression pattern across different stages of tumor growth.

Overall, these findings suggest that as the generations progress, 4T1 cells undergo significant transcriptional shifts under selective pressures, which could favor evolutionary genetic changes that contribute to survival and tumor progression.

### Acclimatory changes and transcriptional trajectory of 4T1 cells

By identifying abundantly expressed genes based on fold change, we observed significant alterations across transcriptomic levels *in vivo*. This uncovered several key changes over time. For instance, transcriptomic expression of various genes changes across different stages when compared to baseline 4T1 cells, suggesting an evolutionary shift of 4T1 cells under *in vivo* environmental pressure (*Fig. 2A*).

**Figure 2:**
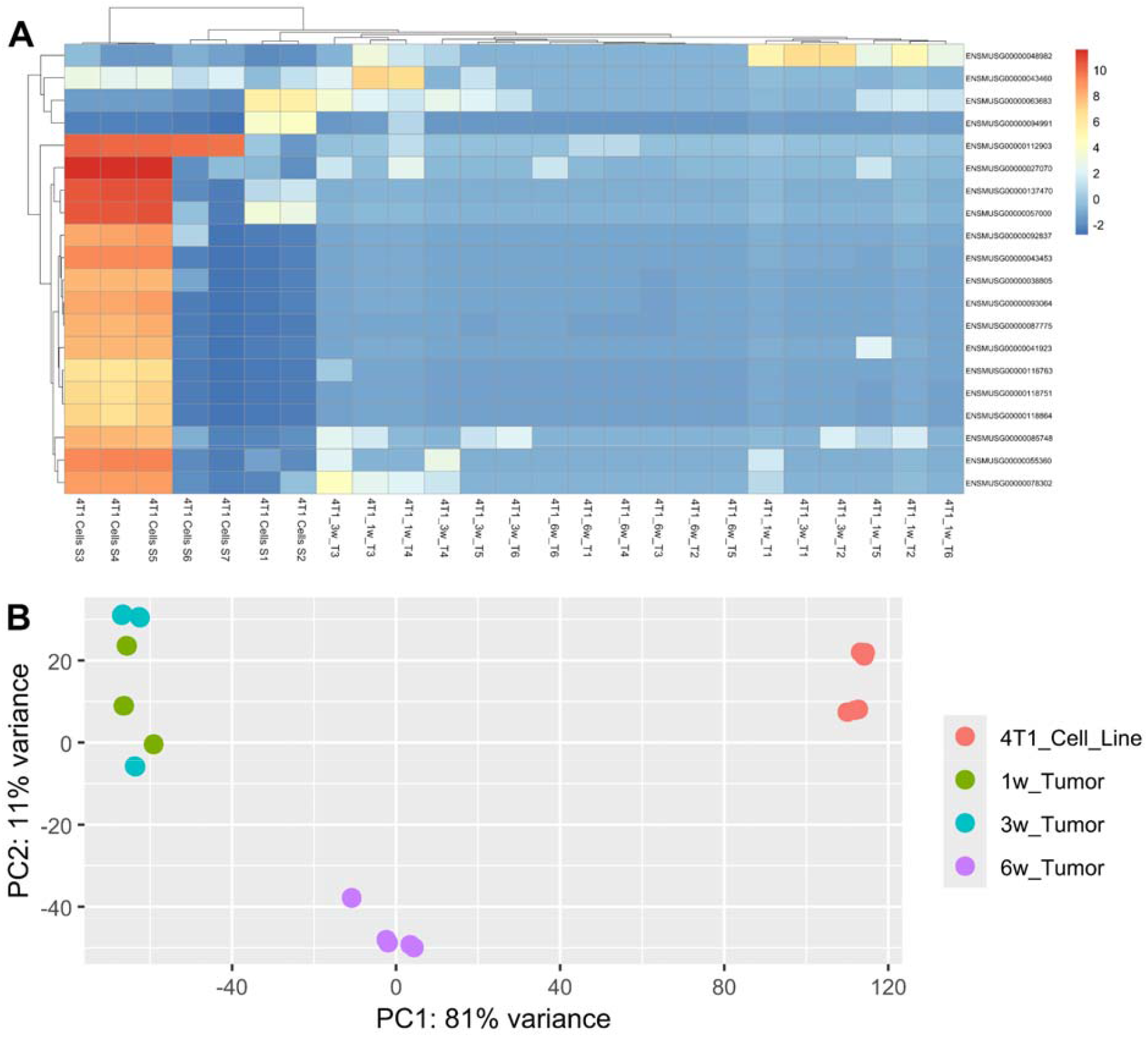
**(A)** Heatmap of highly expressed top 20 genes in 4T1 cells (baseline) and 1-, 3-, and 6-week tumor samples. **(B)** Principal Component Analysis (PCA) for 4T1 cells (baseline) and 1-, 3-, and 6-week tumor samples.

Principal Component Analysis (PCA) also highlights the transcriptional shift in 4T1 cells over time, which provides insights into the conformational flexibility and diversity of conformations that emerged from the trajectory of 4T1 cell growth across different stages (*Fig. 2B*). For instance, 4T1 cell line (red) is separated from all tumor samples along PC1 (81% variance). This indicates a significant transcriptional shift as they acclimate and adapt to a new environment, and in this case, the *in vivo* model system. The tight clustering of 1w and 3w groups indicates similarities in gene expression patterns. As expected, since evolution is a rapid phenomenon and as the higher generation cells (6w) are likely to have more evolutionary changes compared to lower generation cells (1w)^[45]^, we observed a significant separation at 6w samples (*Fig. 2B*). The transcriptional shift from the earlier generation (1w) to the later generation (6w) suggests cumulative evolutionary alterations in key oncogenic pathways related to tumor cell growth^[46]^.

### Gene ontology (GO) analysis: Phenotypic plasticity of gene networks

Hub genes are central nodes in cellular networks, interacting with many other genes, and their interactions can drive tumor cell survival and progression^[47]^. We observed, as indicated earlier, 14,571, 15,036, and 8,925 significantly DEGs (Adj. P ≤ 0.05) at 1w, 3w, and 6w, respectively (*Fig. 1*), which we used for GO analysis. Intersection analysis uncovered 213 common target genes, including ecDNA-associated oncogenes, such as CDK4, WT1 and KIT^[20,48,49]^, which accounted for 0.45% of the total studied genes (*Fig. 3A*). In addition, we identified 27 consistently upregulated genes across all stages (*Fig. 3B*). These genes (illustrated in *Fig. 3A and 3B*) are 4T1 cell-associated genes as they are commonly present across different samples, which we used for further analysis.

**Figure 3:**
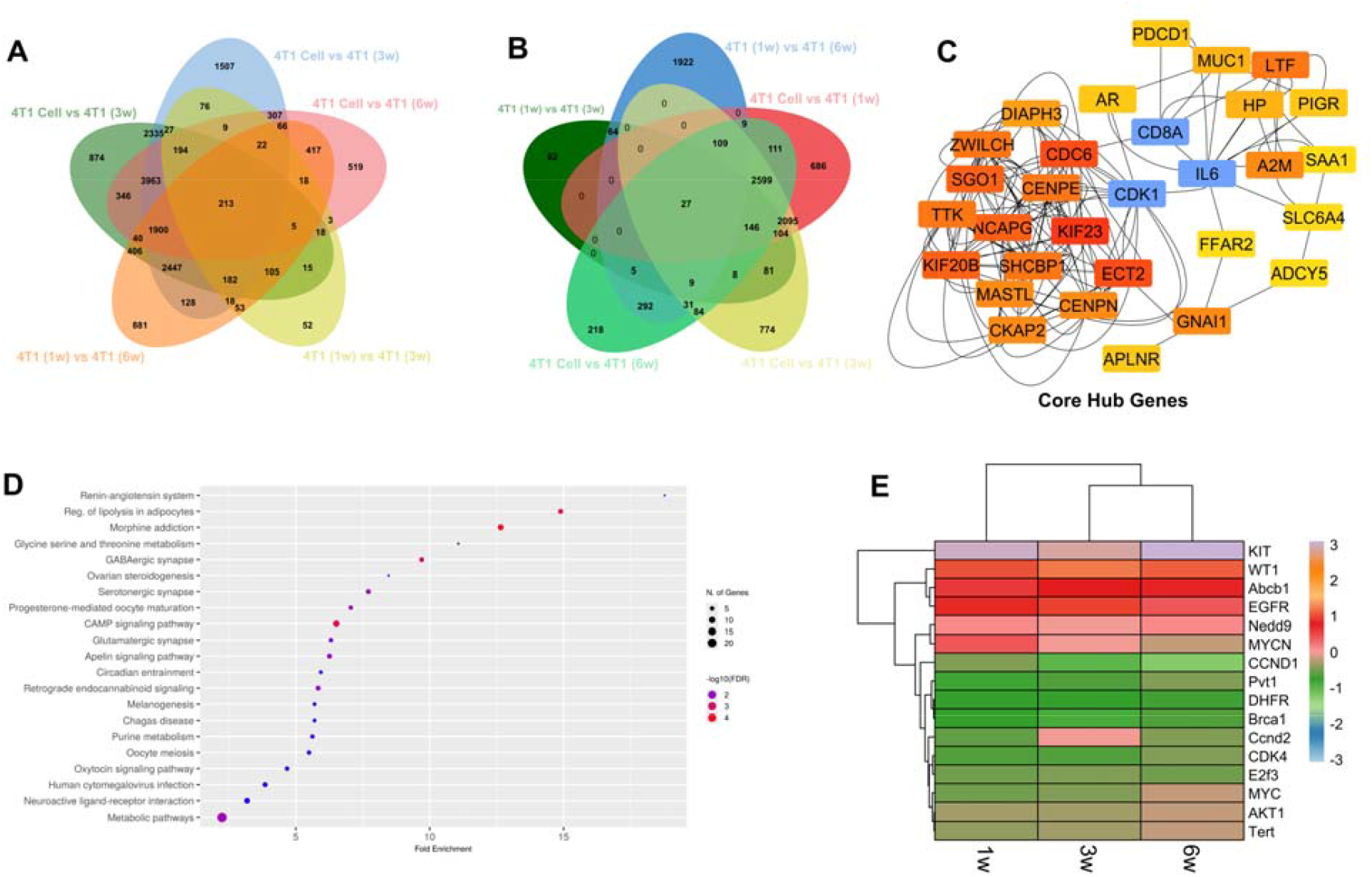
**(A-B)** Venn diagram depicting (A) common genes detected in DEGs and (B) upregulated genes in 4T1 cells and 1w, 3w, and 6w tumor samples. These genes (illustrated in A and B) are 4T1 cell-associated genes as they are commonly present across different samples, which we used for further analysis. **(C)** The highly interconnected hub genes among the DEGs. **(D)** KEGG pathway analysis of the common genes from 4T1 cells. **(E)** Changes in expression for ecDNA-associated genes across different stages of tumor development (1-, 3-, and 6-week post-tumor-implantation).

Interestingly, in late-stage 4T1 tumor samples (i.e., 6w) compared to early-stage 4T1 tumor samples (i.e., 1w), we observed a significantly higher expression of CD8α (a T-cell marker essential for antigen processing and presentation^[50]^), counterbalanced by upregulation of PDCD1 (associated with immunosuppression^[51]^) and CDK1 (associated with tumor aggression^[52]^) (*data not shown*). The identified hub genes, as illustrated in the form of a protein-protein interactions network (*Fig. 3C*), regulate multiple pathways, including signaling, cell cycle regulation, apoptosis, and metabolism (*Fig. 3D*). These findings suggest that as tumor cells undergo evolutionary acclimation and adaptation, they reshape the TME to facilitate their survival and replication.

### Phenotypic plasticity of ecDNA-related genes

Tumor evolution is driven by a complex interplay of genetic and non-genetic factors, including the TME, epigenetic modifications, and, more recently, ecDNA^[19]^. EcDNA facilitates rapid genomic alterations, accelerating tumor adaptation and progression^[19]^. Our investigation identified several oncogenes, including WT1, KIT, EGFR, MYC, and ABCB1, expressed in 4T1 cells (*Table 1*), which were reported by others as ecDNA-associated oncogenes^[20,48,49,53-56]^. These ecDNA genes are known to contribute to treatment resistance^[57,58]^. The transcriptional landscape of these ecDNA-related oncogenes showed dynamic shifts between weeks 1 to 6, reflecting distinct evolutionary phases. For example, early and intermediate stages (1w and 3w) shared similar gene expression patterns, whereas by late-stage (6W), a significant transcriptional reprogramming occurred, leading to the downregulation or loss of certain genes (*Fig. 3E, Table 1*). Interestingly, MYC, a well-known ecDNA oncogene^[20]^, was highly expressed during early-stage tumor acclimation but was progressively downregulated and eventually lost by 6w, suggesting a transition away from MYC dependency (*Fig. 3E, Table 1*). This shift was counterbalanced by the upregulation of WT1, another ecDNA-associated oncogene^[48]^. Additionally, the expression of EGFR, another ecDNA gene^[20]^, was decreased by approximately 50% from 1w and 6w, indicating a potential evolution-driven reduction in EGFR signaling (*Fig. 3E, Table 1*). The drug-resistance gene, ABCB1, remained consistently expressed across different stages, reinforcing its role in sustained chemoresistance^[53]^. Meanwhile, NEDD9, a key regulator of cell migration and invasion^[20]^, followed a distinct transcriptional pattern: initially active, NEDD9 was suppressed at the intermediate stage (3w) and reactivated by 6w (*Fig. 3E, Table 1*). This suggests a period of metastatic dormancy followed by the re-engagement of invasion pathways, aligning with late-stage tumor dissemination. Collectively, these phenotypic shifts suggest that TNBC cells transition through distinct phases: from active proliferation and survival to peak drug resistance, followed by a late-stage metastatic adaptation evident in part by these regulatory changes specific to ecDNA-associated oncogenes. Overall, this highlights the dynamic phenotypic and genomic plasticity of ecDNA-driven tumor progression, where transcriptional reprogramming and ecDNA may enable tumors to evade therapeutic pressures and acquire new survival strategies.

**Table 1.**
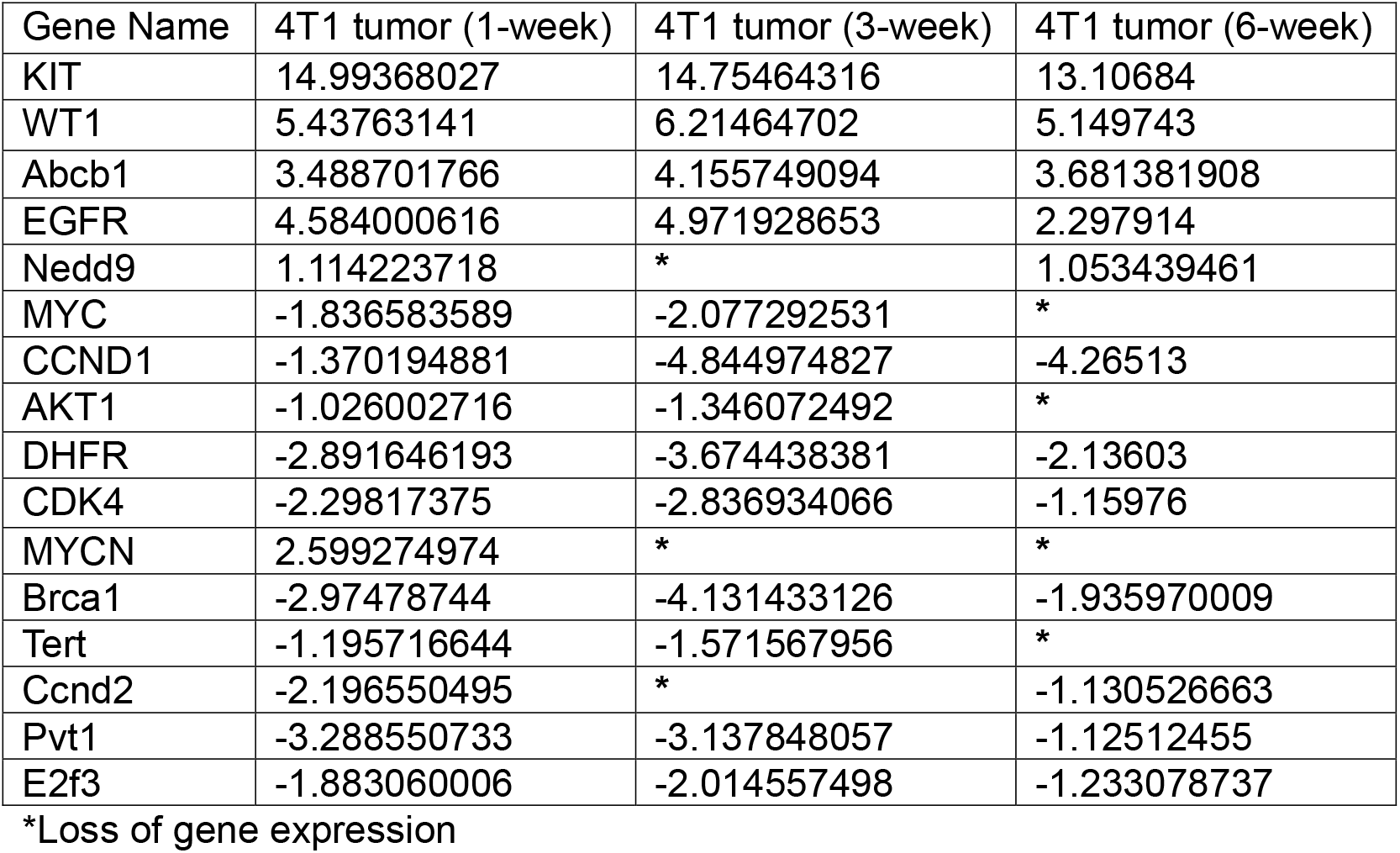
Gene expression level of selected ecDNA genes across different stages of tumor growth (Log2FC)

### Interaction between ecDNA genes and anti-TNBC drugs

Tumor evolution across different stages of tumor growth is likely to induce mutations in various non-chromosomal genes, including ecDNA genes^[21]^. As ecDNAs play a critical role in evolutionary adaptation and drug resistance^[20,21]^, it is expected that mutant ecDNAs (compared to wild-type) are likely to have different types of interaction with anti-TNBC drugs, and that may eventually affect treatment response. However, since the whole-genome sequence (WGS) of datasets used in this study (*Table 1*) was unavailable, we could not evaluate mutations at the ecDNA levels. Bearing this limitation in mind (i.e., missing WGS datasets), we utilized publicly available mutants (which are reflective of evolution-related mutations) of our selected ecDNA genes to evaluate their interactions with DOX or PAC and compared them side-by-side with their wild-type counterparts. Understanding how anti-TNBC drugs interact with ecDNA genes (mutant and wild-type) could shed light on ecDNA-related evolutionary adaptation to anti-TNBC drugs. When hub genes co-evolve with ecDNA genes, they significantly impact drug binding affinity, resulting in differential responses to chemotherapy^[57,58]^. We selected KIT and WT1, two highly expressed ecDNA genes (*Fig. 3E; Table 1*), to investigate their interaction with drugs.

We did not find any report showing DOX interaction with KIT protein. However, in this study, molecular docking analysis showed that wild-type KIT protein has a strong binding affinity with DOX (−8.6 kcal/mol), supported by hydrogen bonds with Leu-595, Cys-673, and Arg-684, along with stabilizing hydrophobic interactions (*Fig. 4A*). However, 14 amino acid residues were mutated in KIT protein, which likely responsible for the reduction of the hydrogen and hydrophobic bonds, resulting in falling the docking score to −7.6 kcal/mol (*Fig. 4B*), which likely plays a role in drug resistance. Conversely, wild-type WT1 protein has a moderate binding affinity to DOX (−6.6 kcal/mol) (*Fig. 4C*), and its mutation (Arg26) introduces new stabilizing interactions with His-373, Arg-372, Arg-394, and Ser-393 that increase the docking score to −7.4 kcal/mol (*Fig. 4D*), which is likely to enhance drug sensitivity. Overall, these findings suggest that anti-TNBC drugs may impose selective pressure when 4T1 cells are grown in the presence of drugs, favoring evolution with the emergence of new clones.

**Figure 4:**
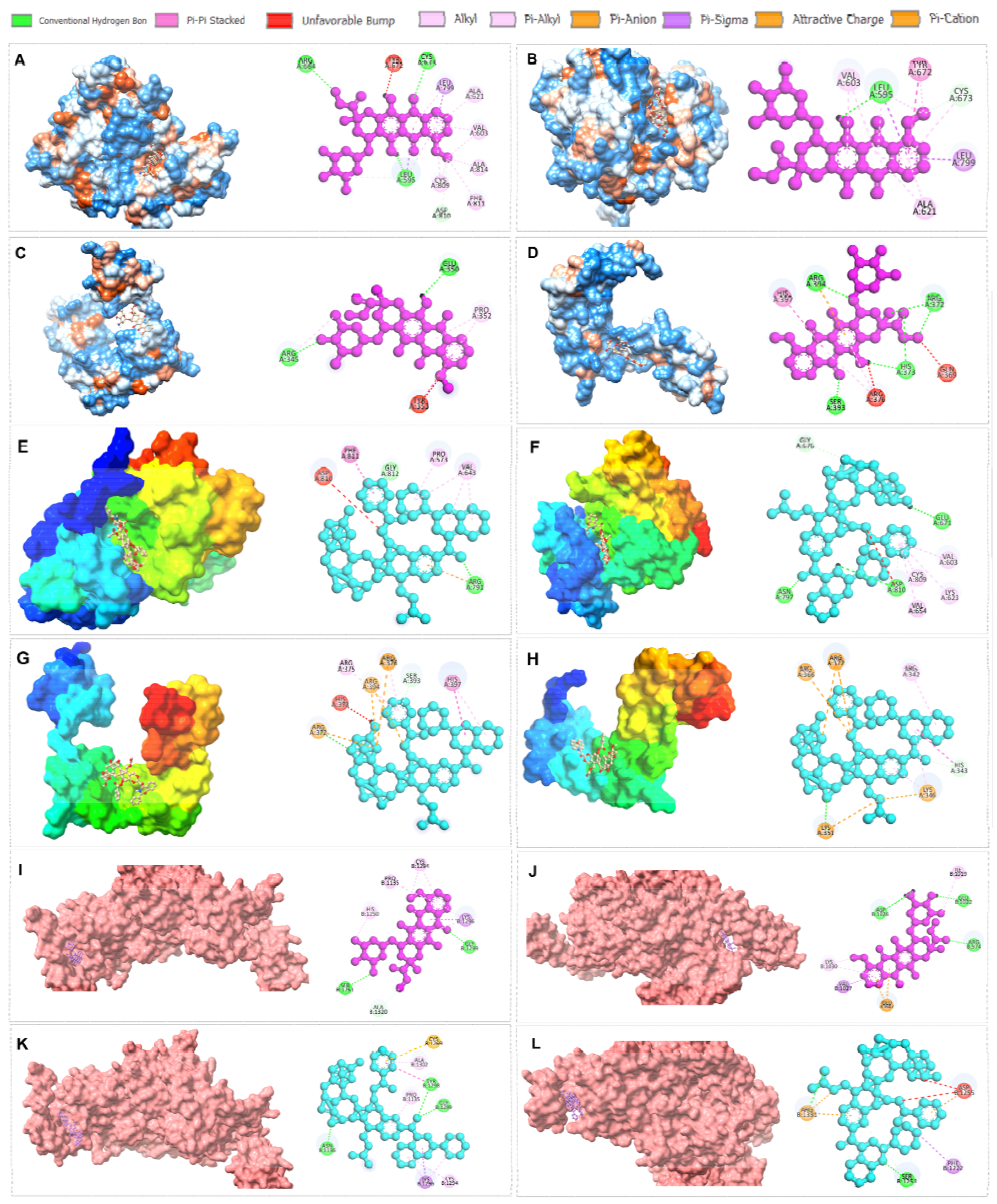
**(A-D)** Doxorubicin interaction with wild-type and mutant ecDNA proteins. (A-B) Doxorubicin interaction with (A) wild-type (PDB: 6MOB) and (B) mutant (PDB: 8PQ9) KIT proteins. (C-D) Doxorubicin interaction with (C) wild-type (PDB: 6B0Q) and (D) mutant (PDB: 6BLW) WT1 proteins. **(E-H)** Paclitaxel interaction with wild-type and mutant ecDNA proteins. (E-F) Paclitaxel interaction with (E) wild-type (PDB: 6MOB) and (F) mutant (PDB: 8PQ9) KIT proteins. (G-H) Paclitaxel interaction with (G) wild-type (PDB: 6B0Q) and (H) mutant (PDB: 6BLW) WT1 proteins. **(I-L)** Drug interaction with wild-type and mutant MSH6 proteins. (I-J) Doxorubicin interaction with (I) wild-type (PDB: 2O8C) and (J) mutant (PDB: 8AG6) MSH6 proteins. (K-L) Paclitaxel interaction with (K) wild-type (PDB: 2O8C) and (L) mutant (PDB: 8AG6) MSH6 proteins.

In response to PAC, wild-type KIT and WT1 proteins demonstrated strong binding affinities (docking scores: −13.1 kcal/mol and −9.9 kcal/mol, respectively), interacting with Arg-791 (KIT) and Arg-372 (WT1) (*Figs. 4E and 4G*). The mutant KIT protein exhibited a greater binding affinity (−15.0 kcal/mol) compared to the wild-type protein, likely due to additional stabilizing hydrophobic interactions with Asn-797, Asp-810, and Glu-671 (*Fig. 4F*). This suggests mutation-related structural shifts. Similarly, WT1 mutation introduced new binding interactions with Lys-351, increasing the docking score to −10.4 kcal/mol (*Fig. 4H*), which may enhance drug sensitivity. These differences in drug-protein interactions illustrate how mutations in ecDNA could directly alter therapeutic responses, either promoting resistance (as with reduced DOX binding affinity to mutant KIT) or potentially enhancing sensitivity (as observed in mutant WT1-PAC interactions). This highlights ecDNA mutations as promising targets for precision drug development.

### Interaction between mutator allele and anti-TNBC drugs

As described above, the selected ecDNA-associated oncogenes undergo significant structural alterations in their binding pockets, weakening drug interactions under predictive selective pressure. Mutations in these genes result in weaker hydrogen bonds, unfavorable steric clashes (bumps), and altered hydrophobic interactions, ultimately reducing drug binding affinity. Additionally, selection pressure may also impact genes associated with genomic instability, such as mutator alleles, which could drive tumor cell evolution and drug resistance^[59]^. Therefore, we investigated DOX or PAC interaction with the common mutator allele, MSH6^[60]^.

Under the predictive selective pressure of DOX, wild-type and mutant MSH6 proteins (G was replaced by E in mutant MSH6 at position 93) show similar binding affinities, approximately −8.3 and −8.6 kcal/mol, respectively (*Figs. 4I and 4J*). However, structural alterations were observed at the drug-binding site in the mutant protein, promoting the formation of three new hydrophobic bonds. This suggests that selective drug pressure could increase the mutation rates, potentially leading to the rapid emergence of beneficial tumor cell adaptations (i.e., drug resistance). However, the binding affinity of PAC for the mutant MSH6 protein (−12.8 kcal/mol) slightly decreased compared to the wild-type protein (−13.5 kcal/mol) and showed one conventional hydrogen bonding interaction with Ser-1251, indicating that the binding pocket of MSH6 was not expanded due to the presence of unfavorable steric clashes (*Figs. 4K and 4L*).

In summary, these studies indicate that 4T1 cells undergo rapid transcriptional shifts across different stages, with changes at the ecDNA levels that could drive tumor cell evolution and drug resistance.

## CONCLUSIONS and IMPLICATIONS

Tumor cells evolve rapidly, which can lead to aggressive tumor growth, the development of invasive and metastatic potential, an increased mutational burden, and, eventually, resistance to treatment^[61,62]^. TNBC, once metastasized, is often lethal and develops resistance to commonly used chemotherapeutics, such as DOX and PAC^[1]^. In this study, using the mouse 4T1 TNBC model system, we examined the transition of tumor cells from *in vitro* to *in vivo* environments. We then further examined changes in tumor cell gene expression at different stages of tumor development under *in vivo* environmental pressures. Rapid tumor cell evolution across generations frequently leads to changes in ecDNA levels and mutations in genes associated with genomic instability (e.g., MSH6)^[12,58]^. Bearing these changes in mind, we also investigated whether mutations in genes associated with ecDNA or genomic instability influence chemotherapy sensitivity or resistance.

During the initial phase, transcriptomic gene expression analysis revealed rapid acclimation of 4T1 cells to the *in vivo* environment since a significant number of genes were differentially expressed at 1-week post-implantation compared to 4T1 cells cultured *in vitro*. This also suggests that acclimatory transcriptional shift favors their growth and survival in response to *in vivo* environment pressures. These pressures likely stem from both internal survival challenges among 4T1 tumor cells and/or external influences from the TME, including the host immune system^[38]^.

It is important to note that 4T cells were implanted in BALB/mice, which are syngeneic to these tumor cells^[63]^. This genetic compatibility facilitates the acclimation and possible later adaptation of 4T1 TNBC cells to the *in vivo* system. Interestingly, while tumor acclimation is generally rapid^[62]^, molecular changes occurred at a slower rate between one and three weeks post-implantation, as indicated by tight clustering in PCA analysis. This slower transcriptional shift can be explained by the fact that 4T1 cells, although syngeneic to BALB/c mice^[63]^, face initial challenges to external pressures arising from the initial immune responses from the host, including infiltration of immune cells into the TME to counteract tumor growth^[63]^.

However, the transcriptional shifts observed after one week suggest that 4T1 cells successfully evade the host immune response by gradually acclimating to the *in vivo* system. Over successive generations, tumor cells often become increasingly adept at immune evasion, as evidenced by a distinct separation in gene expression profiles at six weeks post-implantation compared to earlier time points. This indicates that higher-generation tumor cells undergo more extensive evolutionary changes than lower-generation cells^[45]^, altering pathways critical for tumor cell survival^[46]^.

A key question arises: How do 4T1 cells at earlier generations (e.g., one or three weeks post-implantation) escape the initial external pressure (e.g., host immune response)? A likely explanation is that acclimation favors mechanisms that enable tumor cells to evade immune attack^[64]^. For instance, although CD8α (a marker for CD8^+^ T cells responsible for antitumor immunity^[50]^) was significantly upregulated at six weeks post-implantation, this was counterbalanced by increased expression of immunosuppressive marker PDCD1^[51]^ and tumor-promoting CDK1^[52]^. This suggests that tumor cells evolve mechanisms to suppress immune responses, facilitating their survival and growth in the TME. Thus, it would be interesting to see how 4T1 cells evolve *in vivo* in the absence of PDCD1 using PDCD1-deficient BALB/c mice^[65]^. Further research is needed in this context, especially work that simultaneously compares the genomic composition and transcriptional profiles of the ancestor 4T1 population to replicative 4T1 populations undergoing experimental evolution *in vitro* and *in vivo* in controlled environments.

The transcriptional changes observed at one-, three-, and six-weeks post-implantation (relative to *in vitro* cultured 4T1 cells) include genes from both tumor and non-tumor cells. However, since our focus is on tumor cells (excluding the non-tumor cells, such as immune and other non-tumor cell types residing in the TME), we specifically analyzed for hub genes, which are central nodes in cellular networks and can have primary roles in driving tumor progression^[47]^. The hub genes, commonly expressed in both cultured 4T1 cells and *in vivo* tumor samples, play crucial roles in tumor adaptation to environmental pressures^[66]^. Future research should further investigate key hub genes, particularly those that are highly upregulated, to determine their interactions within tumor cell gene networks and pathways.

One such interaction involves co-amplified hub genes on ecDNA, which are crucial for tumor adaptation and may contribute to drug resistance^[67]^. Although the role of ecDNA in tumor evolution and drug resistance is an emerging field^[19]^, our findings suggest that several ecDNA-associated tumor-promoting genes undergo significant phenotypic changes over time. While some ecDNA-associated oncogenes (e.g., MYC, AKT1, MYCN, EGFR) were significantly downregulated at six weeks post-implantation, others (e.g., KIT and WT1) remained consistently upregulated across different stages, indicating a shift to alternative survival mechanisms. This suggests that tumor progression favors the retention of specific ecDNA genes essential for sustained tumor growth.

Studies are underway in our laboratory to determine how co-amplified hub genes on ecDNA contribute to tumor evolution and resistance to commonly used anti-TNBC drugs, such as anthracyclines (e.g., DOX) and taxanes (e.g., PAC)^[14,15,17]^. Both drugs can induce ecDNA amplifications, potentially influencing drug sensitivity or resistance^[57,58,67,68]^. Importantly, while wild-type ecDNA WT1 or KIT proteins show moderate-to-strong binding affinity to DOX, mutated KIT shows reduced affinity (indicative of drug resistance), whereas mutated WT1 protein shows enhanced binding (indicative of drug sensitivity). Alternatively, mutated WT1 could act as a molecular sink, sequestering DOX away from its site of action (e.g., DNA, TopolI) or impairing its ability to generate reactive oxygen species, which would result in drug resistance^[69]^. In contrast, the binding affinity of PAC increases with mutations in both ecDNA proteins (WT1 or KIT), suggesting enhanced drug sensitivity. These ecDNA protein-drug interaction studies highlight the potential for targeting ecDNA-driven mutations to overcome drug resistance. It remains unclear whether the observed mutant versions of WT1 or KIT persist or are available across different stages of tumor growth since WGS data were unavailable in this study.

Tumor evolution is also associated with increased mutation rates, particularly under environmental pressures that drive adaptive changes and differential therapeutic responses^[70]^. One well-documented example is the emergence of mutations in mutator alleles, such as MSH6, a mismatch repair gene^[12]^. MSH6 mutations increase the mutational burden in cancer cells, potentially leading to chemotherapy resistance^[59]^. Consistent with this concept, our study demonstrates that both drugs interact with mutator alleles, influencing drug response.

Future studies should investigate the evolutionary patterns of ecDNA genes (derived from WGS) over time and their interactions with anti-TNBC drugs. Furthermore, selective drug pressures are expected to drive distinct evolutionary adaptations in extrachromosomal (and chromosomal) genes compared to those observed in drug-free environments. Since our study examined tumor acclimation *in vivo* without drug pressure, future studies will explore tumor cell evolution *in vitro* and *in vivo* under selective drug pressures. In summary, our findings demonstrate that 4T1 cells undergo a significant transcriptional shift from *in vitro* culture to *in vivo* environments, indicative of early tumor cell acclimation. This process continues across different stages of tumor development, with pronounced transcriptional changes observed at later stages, suggesting tumor evolution that may influence differential responses to TNBC therapies.

## Acknowledgments

D.S. was supported in part by a fund from the Division of Academic Affairs at North Carolina A&T State University.

## Author Contributions

M.I.: Performed experiments, data analysis, prepared figures, and wrote the manuscript; P.M.M.: Breast cancer genomics; R.H.N.: Molecular biology and ecDNA-drug interaction; C.J.R.: 4T1 model system; S.H.H.: Bioinformatics; M.T.H.: Overall guidance to the first author; M.D.T.: Evolutionary genomics; J.L.G.: Evolutionary biology, genomics, and overall guidance; DS: Conceptualization, study supervision, data analysis, and wrote the manuscript. All authors edited the manuscript and approved the submission.

