## Supplement File for "Molecular Phenotypic Plasticity Informs Possible Adaptive Change of Triple-Negative Breast Cancer Cells *In Vivo*"

**SUPPLEMENTARY TABLE**

**Table S1:** *NCBI SRA numbers of 4T1 TNBC cells and tumors at different stages of tumor growth (1-, 3-, and 6-week)*

| Serial No. | 4T1 tumor  (1-week) | 4T1 tumor  (3-week) | 4T1 tumor  (6-week) | 4T1 cells |
| --- | --- | --- | --- | --- |
| 1 | SRR10428182 | SRR10428188 | SRR10428194 | SRR12898539 |
| 2 | SRR10428183 | SRR10428189 | SRR10428195 | SRR12898540 |
| 3 | SRR10428184 | SRR10428190 | SRR10428196 | SRR21229974 |
| 4 | SRR10428185 | SRR10428191 | SRR10428197 | SRR21229975 |
| 5 | SRR10428186 | SRR10428192 | SRR10428198 | SRR21229976 |
| 6 | SRR10428187 | SRR10428193 | SRR10428199 | ERR3825656 |
| 7 | - | - | - | ERR3825657 |
